# Exploration of the Roles of HLAs When Predicting Infection Status by T Cell Receptors

**DOI:** 10.1101/2024.11.18.624054

**Authors:** Feng Ding, Si Liu, Wei Sun

## Abstract

T cells are critical components of human immune system. When a cell is infected by a virus, it presents viral peptides on its surface using human leukocyte antigen (HLA) proteins. These peptide-HLA complexes are recognized by T cells through interactions with T cell receptors (TCRs). A human blood sample can contain millions of unique TCRs, which is a sample from the TCR repertoire of the individual. TCR repertoire-wide association studies (TReWAS) aim to evaluate the association between individual TCRs and disease or exposure status. Previous studies have demonstrated that TCRs associated with viral infections can be identified through TReWAS, and such TCRs can be used to predict current or past infection with high accuracy. Notably, many TCRs are strongly associated with specific HLA alleles, suggesting that incorporating HLA information could enhance the accuracy of TReWAS analyses and TCR-based predictions. In our study, we evaluated TCR-based predictions by conditioning on individual HLA alleles or their k-nearest neighbors. We observed improved prediction accuracy for certain HLA alleles. Furthermore, these HLA-specific predictions provide insights into the role of specific HLAs in the infection or disease of interest, offering potential applications in personalized medicine.

## Introduction

The TCR repertoire of an individual, which is defined as the collection of all unique TCRs in that person, captures a historical record of T cell responses. When a T cell recognizes a foreign antigen through its TCR, it undergoes clonal expansion. Following the clearance of antigen-presenting cells, the expanded clone contracts to a stable memory state, where it can persist for years. For example, T cells specific to SARS-CoV-1 (the virus responsible for the 2002 SARS outbreak) have been detected 11 years post-infection [1].

A TCR is composed of an alpha chain and a beta chain, with the beta chain exhibiting greater sequence diversity. A healthy individual typically expresses approximately 10^7^ unique TCR beta chains [2] out of ∼10^14^ possible TCR beta chains [3]. This extreme sequence diversity makes it highly likely that T cells sharing the same TCR arise from the clonal expansion of a single T cell. Consequently, TCRs serve as effective labels for identifying T cell clones.

While most TCRs are unique to an individual, public TCRs shared across individuals have been observed in various immune-related diseases, including infectious diseases (e.g., COVID-19), autoimmune diseases (e.g., type I diabetes), and cancer [4]. Two pioneering studies demonstrated that a TCR Repertoire-Wide Association Studies (TReWAS) can identify infection-associated public TCRs. The burden of these TCRs has been shown to accurately predict current or past infections, such as cytomegalovirus (CMV) [2] or SARS-CoV-2 [5].

A well recognized confounder in TReWAS is the Human Leukocyte Antigen (HLA) composition of each individual [2, 3]. TCRs recognize a protein complex formed by an HLA protein and an antigen. Previous studies have documented clear HLA restrictions for public TCRs and disease-associated TCRs [2, 4–7]. Therefore, disease-associated TCRs can vary with respect to HLA alleles, and accounting for HLA allele information can improve the accuracy of TCR-based predictions. However, to the best of our knowledge, there is no systematic study to evaluate the benefit of accounting for HLA alleles in TReWAS or TCR-based predictions. A potential reason for this gap is the diversity of HLA alleles–there are thousands of HLA alleles and some of them are very rare [8]. Nevertheless, many HLA alleles have relatively high population frequencies, making it feasible to conduct HLA-specific studies for such HLA alleles. In this paper, we aim to predict CMV infection status while accounting for HLA alleles. Our findings reveal that incorporating HLA allele information can improve the accuracy of TCR-based predictions, though the benefits are limited to a subset of HLAs. We also demonstrate that HLA-specific TCR-based prediction of CMV infection can prioritize HLAs that present CMV-derived peptides, highlighting their functional role in protecting against CMV.

## Results

### Accounting for HLA can improve TCR-based prediction

We first demonstrate that a TCR-based predictor conditioned on a specific HLA allele can achieve more accurate predictions of CMV infection status than HLA-agnostic predictors. Using the Emerson data [2], we split the 641 individuals with non-missing CMV status into training (*n* = 513) and test (*n* = 128) sets. We evaluated the accuracy of TCR-based prediction both for all individuals collectively and for subsets of individuals carrying a specific HLA allele. For the latter, we utilized the same training/test split but restricted the analysis to individuals with the specific HLA allele.

Using the training data, we identified CMV-associated TCRs by Fisher’s exact test and selected the top *J* TCRs with p-values below a specified cut-off. Let *x*_*ij*_ indicate whether the *j*-th selected TCR was present in the *i*-th individual. We constructed a predictor for CMV status as 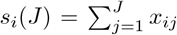. To account for variation in TCR sequencing depth across individuals, we refined the predictor as 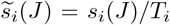, where *T*_*i*_ represents the total number of unique TCRs for the *i*-th individual (Figure 1A-B). The prediction accuracy of 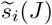 was evaluated on the test data using the area under the ROC curve (AUC), which is appropriate for this dataset given its balanced case/control ratio (∼ 45% CMV+ and ∼55% CMV-individuals). We present the results for several HLAs across different p-value thresholds and compare them with a baseline—a model trained on all training individuals and evaluated on all test individuals. Notably, models trained and evaluated on individuals carrying a specific HLA allele may perform better than the baseline (Figure 1C).

**Figure 1:**
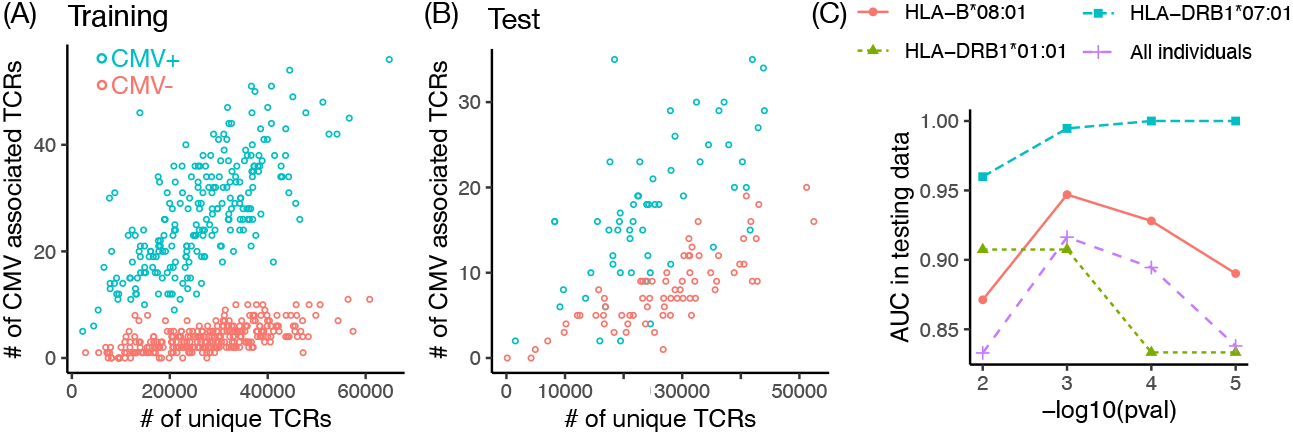
(A-B) The number of unique TCRs versus the number of CMV-associated TCRs (p-value < 10−3) per individual in the training and test data, respectively. (C) The AUC of CMV status prediction in test data, trained and evaluated either across all individuals or specifically for those carrying a particular HLA allele. Each predictor is based on CMV-associated TCRs identified at varying p-value cutoffs (X-axis). The total sample sizes for individuals carrying HLA-B08:01, HLA-DRB101:01, and HLA-DRB1*07:01 are 138, 88, and 141, respectively, with approximately 80% allocated to training and 20% to testing.

### A systematic study of HLA-specific predictions

In the previous section, we compared HLA-specific AUCs to baseline AUCs computed using all test individuals. In this and the following sections, we will compare the HLA-specific AUCs versus a different baseline: while the baseline model is still trained on all training individuals, the AUCs are evaluated only on test individuals who carry a specific HLA allele.

There are two classes of HLAs: HLA-I and HLA-II. HLA-I proteins presents peptides of 8-11 amino acids (AAs) derived from intracellular proteins to CD8+ T cells (cytotoxic T cells). HLA-II proteins present peptides of 12-20 AAs derived from extracellular proteins to CD4+ T cells (T helper cells). An HLA-I protein is encoded by one of three highly polymorphic genes: HLA-A, HLA-B, HLA-C [9]. Most HLA-II proteins are encoded by one of three gene pairs: HLA-DRA/HLA-DRB1, HLA-DPA1/HLA-DPB1, and HLA-DQA1/HLA-DQB1 [10]. Except for HLA-DRA, all the other HLA-II genes are highly polymorphic. Since each gene’s maternal and paternal copies may be different, each human has up to 6 different HLA-I alleles (2 for each of HLA-A, B, and C genes), and up to 10 HLA-II alleles (2 for DR, and 4 for DP and DQ). Many HLA-I or HLA-II alleles are very rare in the human population, making it infeasible to study HLA-specific TCR-disease associations. In this study, we focus on the HLAs that are present in at least 40 individuals.

For an HLA-specific TReWAS, we selected TCRs associated with CMV by restricting the analysis to individuals carrying a specific HLA allele. TCRs with p-values smaller than 0.001 were then used to predict CMV status, with prediction accuracy evaluated by AUC. We chose p-value 0.001 as default since the baseline model achieves highest performance at this p-value cutoff (Figure 1(C)). See Methods section for details of the association analysis and the prediction model. For comparison, we also calculated HLA-agnostic AUCs. In this case, TCRs associated with CMV were identified from all training individuals, regardless of HLA alleles, and the resulting prediction model was evaluated among individuals carrying specific HLA alleles.

Our results showed that the HLA-specific AUC was higher than the HLA-agnostic AUC for several HLA alleles. These included one HLA-I allele (HLA-C07:02) and five HLA-II alleles: HLA-DRB107:01, HLA-DRB103:01, HLA-DQA01:01-DQB05:01, HLA-DQA02:01-DQB02:02, and HLA-DQA05:01-DQB*02:01 (Figure 2(A), Supplementary Table 1).

**Figure 2:**
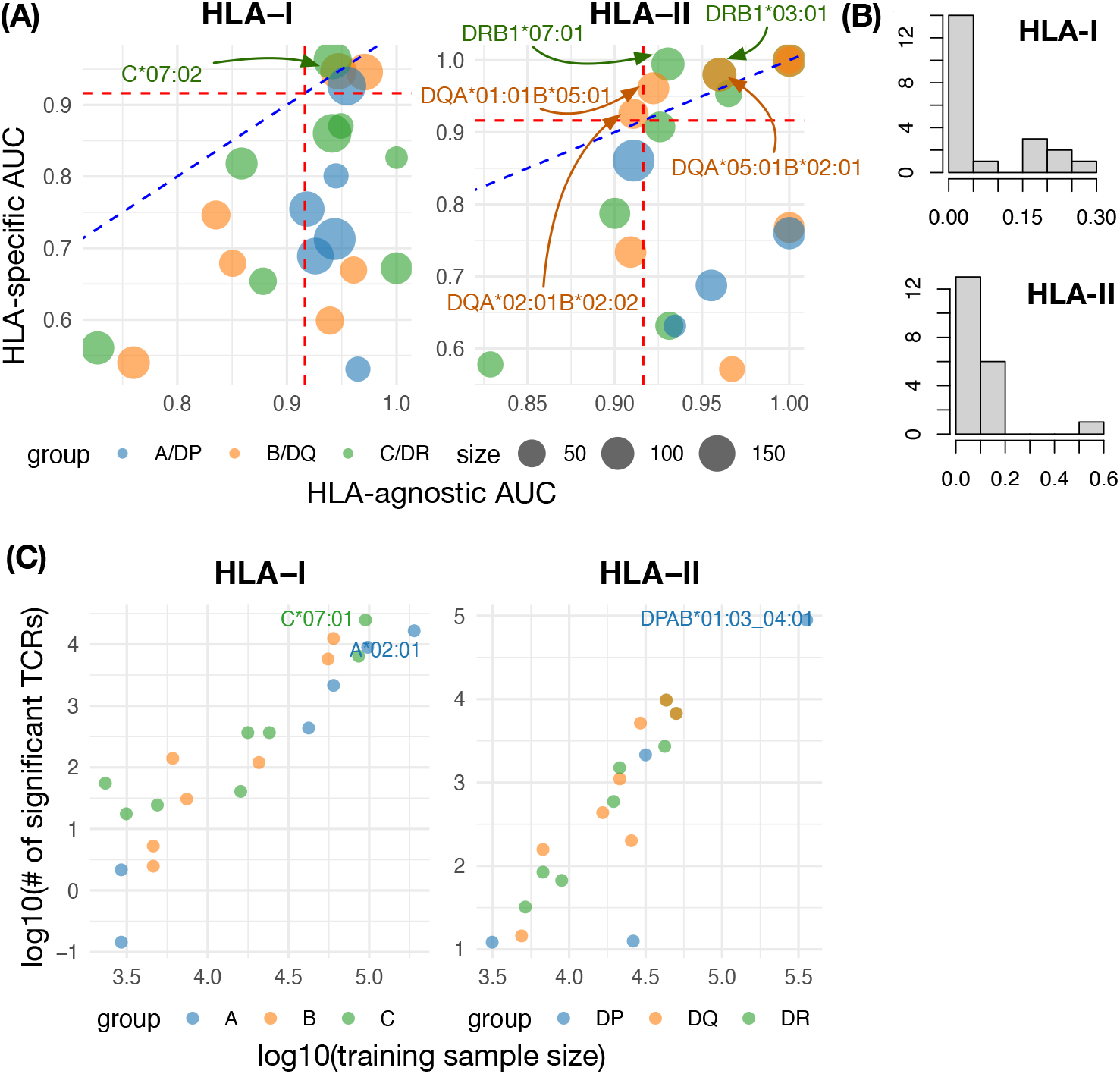
(A) The HLA-agnostic AUC versus HLA-specific AUC for each HLA allele. The red broken lines indicate the AUC when applying HLA-agnostic prediction model to all test individuals. (B) The ratio of the the number of CMV-associated TCRs for each HLA (Y-axis) versus the 280 CMV-associated TCRs from HLA-agnostic analysis. (C) The relation between training data sample size and the number of CMV-associated TCRs across HLA alleles.

However, for most HLA alleles, the HLA-specific AUC was lower than the HLA-agnostic AUC. We hypothesize that this could be due to the limited relevance of certain HLAs to CMV or the smaller number of HLA-specific CMV-associated TCRs identified, likely due to limited sample size. To investigate these hypotheses, we calculated the number of HLA-specific CMV-associated TCRs for each HLA allele and compared them to the 280 HLA-agnostic CMV-associated TCRs. In most cases, the number of HLA-specific CMV-associated TCRs is less than 30% of the HLA-agnostic CMV-associated TCRs (Figure 2(B)). The number of HLA-specific CMV-associated TCRs is strongly associated with sample size (i.e., the number of individuals in the training set carrying a specific HLA allele) (Figure 2(C)). This suggests that increasing the training set size for specific HLA alleles would likely result in more significant TCRs and, consequently, improved prediction accuracy.

### Combining HLA-specific TCRs and HLA-agnostic TCRs to make prediction

Previous analyses demonstrated that the HLA-agnostic model performs well for most HLA alleles. This is likely because the HLA-agnostic model was trained on larger sample sizes, enabling it to identify TCRs with weaker but consistent TCR-CMV associations across multiple HLAs. In contrast, the HLA-specific model may capture stronger associations that are unique to specific HLAs. To leverage the strengths of both approaches, we combined the TCRs identified by the HLA-agnostic and HLA-specific models into a single predictive framework, referred to as the Combined model.

Our results show that the combined AUC is higher than the HLA-specific AUC for nearly all HLA-I alleles. Additionally, for a few HLA-I alleles, the combined AUC exceeds the HLA-agnostic AUC (Figure 3(A)). For HLA-II alleles, the combined model not only outperforms the majority of HLA-specific models but also achieves higher AUCs than the HLA-agnostic model for many HLAs (Figure 3(B)). Overall, the combined AUCs are similar or higher than the HLA-agnostic AUCs (Supplementary Table 1).

**Figure 3:**
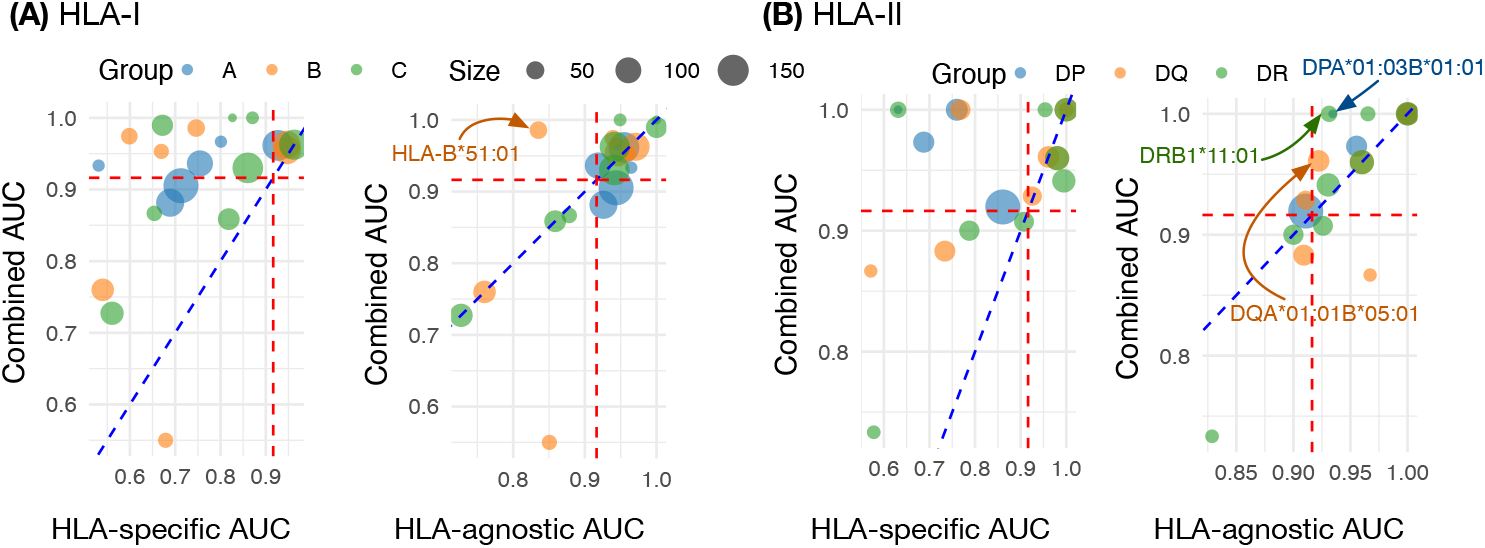
Combined AUC versus the HLA-specific and HLA-agnostic AUC for HLA-I and HLA-II alleles respectively.

### HLA-specific prediction can identify CMV-associated HLAs

HLA-specific prediction of infection status can provide insights into HLAs that are functionally associated with infection. If an HLA allele presents multiple peptides from a pathogen, the TCR-based predictor for that HLA is expected to demonstrate relatively high performance. As a baseline, we also assessed the co-occurrence of HLA alleles and CMV infection using Fisher’s Exact Test. However, this method did not identify any significant associations after correcting for multiple testing (Figure 4(A)).

**Figure 4:**
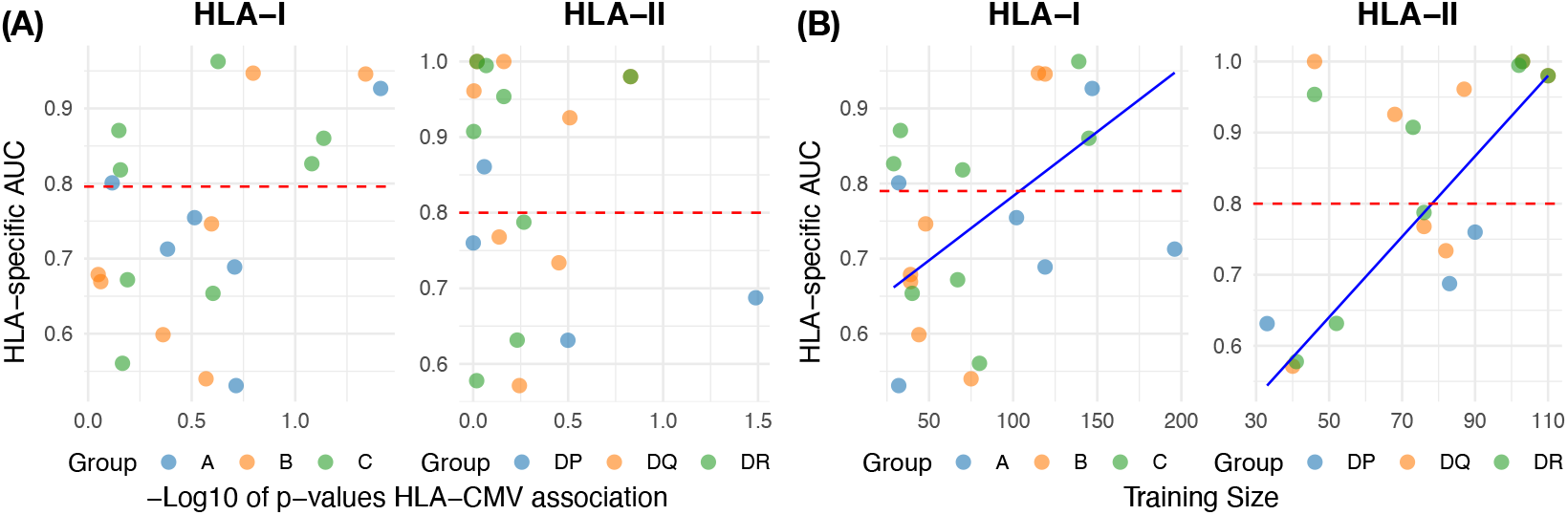
(A) The HLA-Specific AUC (Y-axis) versus the p-values of HLA-CMV association (co-occurrence by Fisher’s exact test) on -Log10 scale (X-axis), for HLA-I and HLA-II respectively. The horizontal line at 0.8 is just for visualization purpose. (B) The HLA-Specific AUC (Y-axis) versus the Training Size (X-axis), for HLA-I and HLA-II respectively. An outlier HLA-DPA*01:03DPB:04*01 that has very large training sample size is removed. See Supplementary Figure 1 for the plot with this outlier.

The HLA-specific AUC varies from 0.5 to 1. While higher AUC values suggest that the corresponding HLA may present more CMV peptides, it is not clear how to choose a cutoff for AUC (Figure 4(A)). In addition, the AUC is positively associated with training sample size. Thus we fit a regression model for the relation between AUC and training sample size. Those HLAs above the regression line have higher AUCs than expected based on sample size, and thus are more likely to present CMV peptides (Figure 4(B), Supplementary Table 1).

### Borrowing information across HLA alleles

To address the challenge of limited sample sizes for individual HLA alleles, we implemented a strategy that borrows information across similar HLA alleles. For each HLA allele, we identified its k-nearest neighbors (k-NN) using a sequence similarity metric based on BLOSUM62 scores. By default, we selected up to four nearest neighbors with similarity scores greater than 0.9. Individuals carrying any of these HLA alleles were combined into a single training set to identify CMV-associated TCRs using a weighted logistic regression. The weights were determined by the similarity of an HLA allele to the HLA of interest. See Methods section for details. Model performance was then evaluated on test data, using only individuals carrying the HLA allele of interest, based on AUCs. We refer to this approach as the KNN model.

The KNN model demonstrated improved prediction accuracy compared to HLA-specific models for many relatively rare HLA alleles. However, it outperformed the HLA-agnostic model for only a small subset of HLA alleles, specifically HLA-B07:02, HLA-C06:02, HLA-DRB101:01, HLA-DRB107:01, HLA-DRB113:05, and HLA-DQAB01:02 06:04 (Supplementary Figures 2-3, Supplementary Table 1).

## Discussions

TCRs associated with pathogen infections can be identified through a TReWAS. These TCRs can then be used to predict infection status with high accuracy. A TCR-based predictor is conceptually analogous to a polygenic risk score (PRS), which predicts the risk of a complex trait using the weighted summation of single nucleotide polymorphism (SNP) genotypes [11, 12]. However, there are two key differences between the two problems. First, while most SNPs have relatively weak associations with diseases (e.g., odds ratios typically range from 1 to 1.3, with a few SNPs showing strong effects), the association strength between a TCR and a disease can vary widely, with odds ratios ranging from 1 to 10 or higher (Supplementary Figure 4). Second, SNP genotypes often exhibit strong correlations due to linkage disequilibrium (LD), which must be accounted for in PRS calculations [13]. In contrast, TCRs generally show no strong correlations, except for their associations with specific HLAs [4].

The focus of this paper is accounting for HLA effects in TReWAS and TCR-based predictions. We investigated whether conditioning on HLAs could improve the accuracy of predicting CMV infection status using TCRs. Our results indicate that HLA conditioning improved prediction accuracy for a few HLA alleles. A likely reason that HLA-specific prediction models often underperform compared to HLA-agnostic models is the limited sample size available for training the former. Borrowing information across HLA alleles can partially mitigate this limitation but is effective for only a small number of HLAs in this CMV data. In this study, we evaluated HLA similarities using the amino acid sequence similarity scores from the BLOSUM62 matrix. However, there may be more effective methods to quantify HLA similarities.

A promising approach proposed in this paper is the combined model, which integrates HLA-specific and HLA-agnostic CMV-associated TCRs to improve prediction accuracy. This combined model also provides a foundation for more personalized predictions of infection status based on an individual’s unique HLA profile. Specifically, HLA-specific TCRs corresponding to all of an individual’s HLA alleles can be aggregated with HLA-agnostic TCRs to generate a personalized prediction. However, achieving optimal results may require assigning appropriate weights to different HLAs, which is a challenge that warrants further studies, preferably for studies with larger sample sizes.

While our focus has been on predicting CMV infection status, the methods developed in this work are generalizable and can be applied to other immune-related diseases and conditions.

## Methods

### Data

We analyzed the TCR and HLA data of 666 individuals, as reported by Emerson et al. [2] and Dewitt et al. [4]. The HLA dataset included presence/absence of 215 HLA alleles, with some missing values. We selected 1, 098, 738 TCRs that were observed in at least seven individuals for our analysis. The CMV infection status of the 666 individuals were reported by Emerson et al. [2]. After excluding individuals with missing CMV infection status, our final dataset consisted of 641 individuals.

### HLA-agnostic model

We divided the 641 individuals into training and test sets using an 80/20 split, resulting in 512 individuals in the training set and 129 in the test set. Within the training set, we conducted a TCR-CMV association analysis for each TCR using Fisher’s Exact Test. By default, any TCR with a p-value smaller than *α* = 0.001 was classified as CMV relevant. To ensure the robustness of our findings, we also evaluated results across different p-value cutoffs to verify that they were not overly sensitive to the chosen threshold (Figure 1(C)). At the default p-value cutoff of 0.001, the number of significant TCRs (*J*) was 280.

For the *i*-th individual in the test set, we calculated a predictor 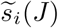, which represents a read-depth-corrected burden score for the CMV-associated TCRs identified during training. This predictor is referred to as “HLA-agnostic” because HLA information was not incorporated during the training phase. We assessed the prediction performance of 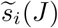 in various subsets of the test set (i.e., individuals carrying specific HLA alleles) by AUC of the ROC curves.

### HLA-specific model

For each HLA allele, we employed a training scheme similar to the HLA-agnostic model, with one key difference: restricting the analysis to training individuals carrying the specific HLA allele of interest. We evaluated HLA-specific models using the same test sets as the HLA-agnostic model, i.e., the test individuals carrying the specific HLA allele.

Because some HLA alleles are relatively rare, the corresponding HLA-specific training sample sizes can be very small, potentially leading to unstable results. To address this, we pre-filtered HLA alleles and included only those carried by at least 40 individuals. For alleles present in 40 to 70 individuals, we further improved model robustness by aggregating results across 10 randomized training/testing splits. Specifically, the dataset was split 10 times using the same 80/20 ratio, and the analysis was conducted independently for each split. The AUCs from the test data were then averaged to produce the final results.

### Combined model

In the combined model for an HLA allele, we integrated TCRs associated with CMV identified by both the HLA-agnostic model and the corresponding HLA-specific model. The HLA-agnostic model selected 280 TCRs using a p-value cutoff of 0.001. For each HLA allele of interest, we identified additional CMV-associated TCRs using the same p-value cutoff of 0.001 in the HLA-specific model.

During the testing phase, we evaluated the accuracy of CMV infection status prediction for individuals carrying the specific HLA allele. For each allele, the TCRs identified by the HLA-agnostic and HLA-specific models were pooled. Using this combined set of TCRs, the predictor 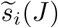 was calculated as in previous HLA-agnostic analyses.

### Borrow information across HLAs

The similarity among HLA alleles was calculated based on the amino acids in the HLA pseudo sequences, which are the residues likely to interact with TCRs or peptides. Specifically, pseudo sequences are 40 amino acids long for HLA-I alleles and 45 amino acids long for HLA-II alleles [14]. Similarity calculations were conducted separately for HLA-I and HLA-II alleles. For any two HLA alleles *a*_1_ and *a*_2_, the similarity score was computed using the formula

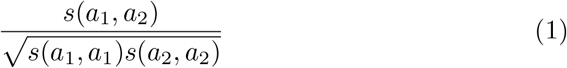

following the approach described by [15], where *s*(*a*_1_, *a*_2_) was the summation of BLOSUM62 matrix values for aligning the amino acids of *a*_1_ and *a*_2_.

For each HLA allele, we calculated its similarity to all the other HLAs and identified its neighbors, defined as those with a similarity score greater than 0.9. If there were more than four such neighbors, the top four nearest neighbors were selected. We performed TReWAS for each HLA alleles using individuals who carried the target HLA allele or one of its nearest neighbors. To account for variation in similarity scores, so that closer neighbors contributed more, we used weighted logistic regression instead of Fisher’s exact test for the TReWAS analysis.

During the training phase, we first log-transformed the total number of TCRs carried by each individual, denoted as log(*rd*), where *rd* represents TCR readdepth. For each TCR, we then fit the following logistic regression models:

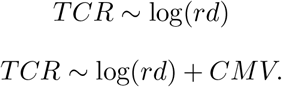

where both *TCR* and *CMV* were binary indicators denoting whether an individual carried a specific TCR or had contracted CMV, respectively. We used TCRs as the response variable to account for the effect of TCR read-depth. After fitting the models, we examined the slope parameter for *CMV*, and retained only TCRs with positive slopes, as we were interested in TCRs co-occurring with CMV. The association between *TCR* and *CMV* was then assessed using a likelihood ratio test comparing these two models. By default, the top *J* significant TCRs were selected using a p-value cutoff of 0.001. For each individual in the test set, we calculated the predictor 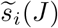, and we referred to this model as the KNN model.

To evaluate the performance of the KNN model, we tested it on the same test sets as those used for the HLA-agnostic or HLA-specific models, i.e, individuals carrying the specific HLA allele.

## Supporting information

Supplemental Figuers

