## Supplemental Figuers for "Exploration of the Roles of HLAs When Predicting Infection Status by T Cell Receptors"

### Supplementary Materials for "Exploration of the Roles of HLAs When Predicting Infection Status by T Cell Receptors"

November 19, 2024

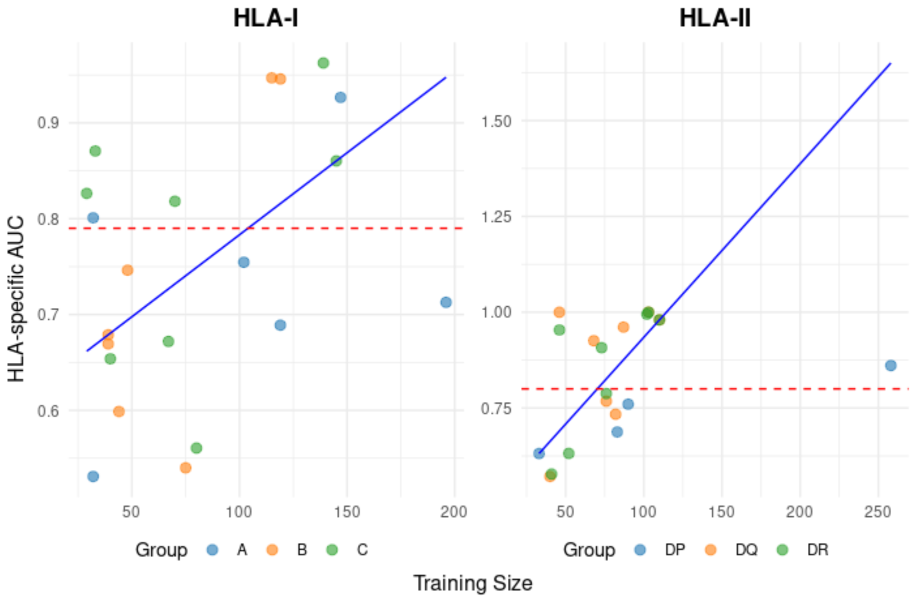


Figure 1: The HLA-Specific AUC (Y-axis) versus the Training Size (X-axis), for HLA-I and HLA-II respectively(with the outlier HLA-DPA*01:03-DPB:04*01 included).

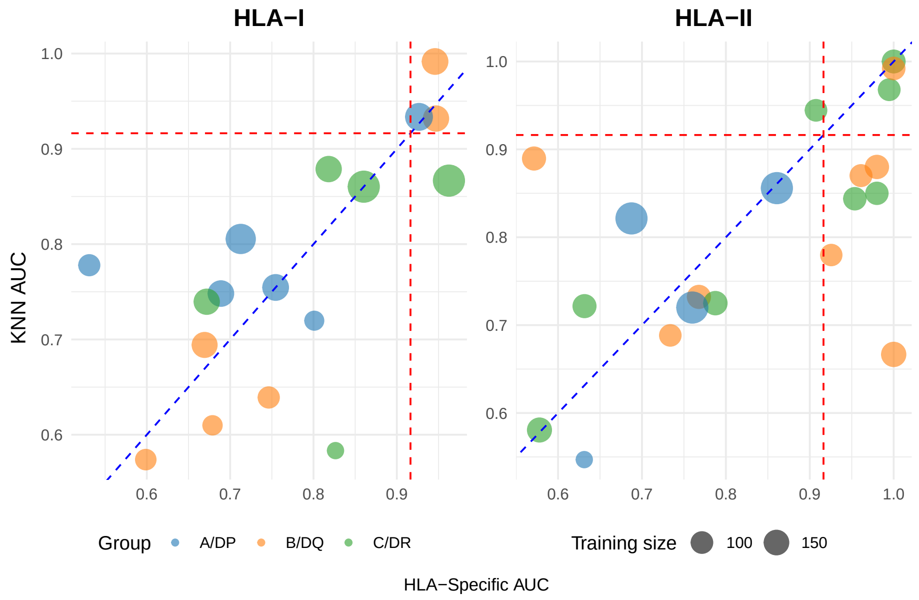


Figure 2: Comparison of AUCs for the prediction of CMV infection status using HLA-specific model or K-nearest neighbor (KNN) model for HLA-I or HLA-II alleles.

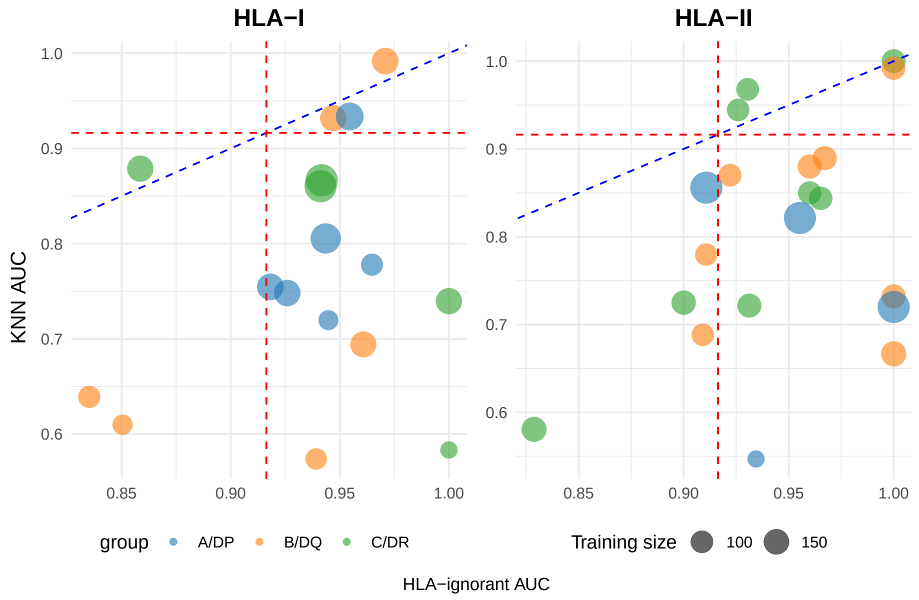


Figure 3: Comparison of AUCs for the prediction of CMV infection status using HLA-ignorant model or K-nearest neighbor (KNN) model for HLA-I or HLA-II alleles.

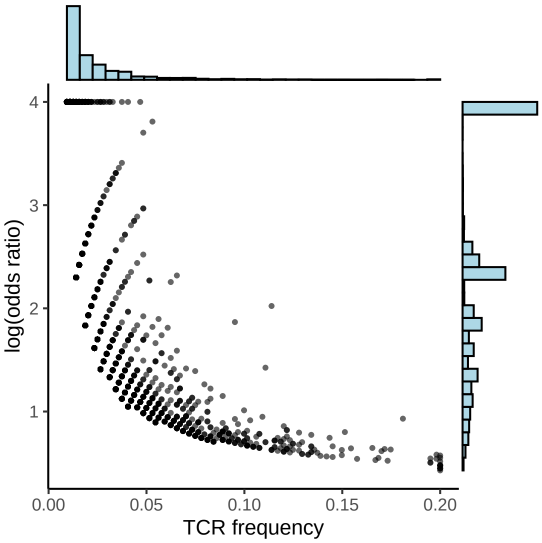


Figure 4: The population frequency (X-axis) and log odds ratio estimates (Y-axis) for 4,868 TCRs whose association p-values with CMV status are smaller than 0.01 using Emerson data [1]. The TCR frequency is truncated at 0.2 for visibility. Some TCRs' odds ratios are infinity (i.e., they only appear in CMV+ individuals but not CMV-individuals), and such infinite log odds ratios are set to be 4.
